# Fatty sweet symphony: Decoding distinct ganglioside patterns of native and differentiated mesenchymal stem cells by a novel glycolipidomics profiling strategy

**DOI:** 10.1101/2022.04.11.487866

**Authors:** Katharina Hohenwallner, Nina Troppmair, Lisa Panzenboeck, Cornelia Kasper, Yasin El Abiead, Gunda Koellensperger, Leonida M. Lamp, Jürgen Hartler, Dominik Egger, Evelyn Rampler

**Affiliations:** Department of Analytical Chemistry, Faculty of Chemistry, University of Vienna, Vienna, Austria; Vienna Doctoral School in Chemistry (DoSChem), University of Vienna, Vienna, Austria; Institute of Cell and Tissue Culture Technologies, University of Natural Resources and Life Sciences, Vienna, Austria; Institute of Pharmaceutical Sciences, University of Graz, Graz, Austria; Field of Excellence BioHealth, University of Graz, Graz, Austria

**Keywords:** Ganglioside, mesenchymal stem cells, differentiation, human, glycolipidomics, mass spectrometry, LC-MS^n^, automated annotation

## Abstract

Gangliosides are an indispensable glycolipid class concentrated on cell surfaces with a critical role in stem cell differentiation. Nonetheless, owing to the lack of suitable methods for scalable analysis covering the full scope of ganglioside molecular diversity, their mechanistic properties in signaling and differentiation remain undiscovered to a large extent. This work introduces a sensitive and comprehensive ganglioside assay based on liquid chromatography, high-resolution mass spectrometry, and multistage fragmentation. Complemented by an open-source data evaluation workflow, we provide automated in-depth lipid species-level and molecular species-level annotation based on decision rule sets for all major ganglioside classes. Compared to conventional state-of-the-art methods, the presented ganglioside assay offers (1) increased sensitivity, (2) superior structural elucidation, and (3) the possibility to detect novel ganglioside species. A major reason for the highly improved sensitivity is the optimized spectral readout based on the unique capability of two parallelizable mass analyzers for multistage fragmentation. In addition to the significant technological advance, we identified 263 ganglioside species including cell-state-specific markers and previously unreported gangliosides in native and differentiated human mesenchymal stem cells. A general increase of the ganglioside numbers upon differentiation was observed as well as cell-state-specific clustering based on the ganglioside species patterns. By proving the predictive power of gangliosides as ubiquitous cell state-specific markers, we demonstrated the high throughput universal capability of our novel analytical strategy, which comes with new insights on the biological role of gangliosides in stem cell differentiation. Our analytical workflow will pave the way for new ganglioside- and glycolipid-based clusters of differentiation markers to determine stem cell phenotypes.

## Introduction

### The biological role of gangliosides

Gangliosides play a crucial structural role in the curvature of the plasma membrane. They are involved in many critical biological pathways related to cell-cell communication, cellular growth, host-pathogen interaction, and signal transduction. Gangliosides protrude from eukaryotic cell surfaces. Their mobile hydrophilic glycan moiety is presented to the outside, and the lipid moiety is cohesively integrated into the hydrophobic plasma membrane. The glycan moiety of gangliosides belongs to the glycocalyx, a dense gel-like matrix surrounding the plasma membrane of a cell, also known as “sweet husk”, which is active in various cellular processes(Möckl, 2020). Gangliosides can serve as ligands and modulate the activity of membrane proteins depending on both the oligosaccharide head group and the ceramide anchor. They belong to the group of acidic glycosphingolipids containing at least one sialic acid.

The highest concentrations of gangliosides are found in the central nervous system(Yu et al., 2012), and they are involved in several memory-related diseases, including Alzheimer’s, Parkinson’s, Huntington’s disease, AIDS-related dementia, or cancer(Sipione et al., 2020). Recently, it was observed that gangliosides facilitate viral entry of SARS-CoV-2, highlighting their crucial interaction role within the plasma membrane(Nguyen et al., 2022). In addition to their essential disease-related functions, ganglioside patterns vary with development stage and age. Dramatic changes in expression levels of gangliosides were observed during neurodevelopment(Ica et al., 2020; Yu et al., 2012). Ranging from the expression of simple gangliosides with low numbers of sugars attached, e.g., GM3 and GD2 in early stages, to more complex gangliosides with higher sugar content in later developmental stages, particularly GM1, GD1, and GT1(Yu et al., 2012). The strong influence of ganglioside composition in neurodevelopment was also expected in stem cell development. This hypothesis triggered investigations directed towards ganglioside biomarkers in differentiation processes(Kwak et al., 2006; Moussavou et al., 2013). During human embryonic stem cell differentiation, a switch in the core structures of glycosphingolipids globo- and lacto- to ganglio-series was observed(Liang et al., 2010), leading to distinct alterations of specific glycosphingolipids(Bergante et al., 2014).

Mesenchymal stem/stromal cells (MSCs) comprise a heterogeneous cell population of non-hematopoietic stem cells, which are prevalently isolated from adipose tissue(Hattori et al., 2004), bone-marrow(Conget and Minguell, 1999), or birth-associated tissues and fluids(Moretti et al., 2009; Witkowska-Zimny and Wrobel, 2011). Due to their high proliferation and differentiation capacity, immunomodulatory effects on the innate and adaptive immune system(Müller et al., 2021), as well as anti-inflammatory and trophic effects on neighboring cells(Fu et al., 2017), are observed. They represent outstanding candidates for cell-based therapies and regenerative medicine applications. As a single marker for the characterization of MSCs was not discovered yet, the International Society for Gene and Cellular Therapy (ISCT) defined minimum criteria for the characterization of MSCs: (1) plastic-adherence, (2) differentiation into adipogenic, chondrogenic, and osteogenic lineage, and (3) positive expression of cluster of differentiation (CD) markers including CD90, CD73, and CD105, and lack of expression of CD14, CD19, CD34, CD45, and HLA-DR(Dominici et al., 2006). To enhance the quality control of cell-based therapy products, additional indicators other than the surface profile expression and differentiation capacity are required to monitor and define the stem cell phenotype during *ex vivo* culture. Gangliosides have been discussed as stem cell and lineage-specific differentiation markers before. For example, GM1, GM3, and GD2 were expressed in umbilical cord-and bone marrow-derived MSCs(Freund et al., 2010; Jin et al., 2010; Martinez et al., 2007). Furthermore, gangliosides were differentially expressed during neural(Kwak et al., 2006) and osteogenic(Bergante et al., 2014, 2018; Moussavou et al., 2013) differentiation of MSCs. In accordance with these earlier studies, we recently found gangliosides to be upregulated in adipocytes compared to their human mesenchymal stem cell (MSC) progenitors(Rampler et al., 2019). However, a comprehensive analysis of the expression pattern of gangliosides in native and differentiated MSCs is missing, although gangliosides show huge structural variations in the glycan and lipid part depending on the cell type and state(Sarbu et al., 2016; Sonnino and Chigorno, 2000). As the ceramide structure and number of sialic acids or other sugar parts lead to changed properties, these structural fluctuations influence the membrane surrounding glycocalyx as well as cell signaling and development cascades. Gangliosides expressed in pluripotent, multipotent, and cancer stem cells have been traditionally identified by biochemical and immunological analysis(Jin et al., 2010; Kwak et al., 2006; Yanagisawa, 2011). Although several fundamental studies indicated that gangliosides could be markers either for cell lineage, cell state, or function in different biological contexts(Jin et al., 2010; Moussavou et al., 2013; Ngamukote et al., 2007), utilization of this observation failed so far, in part due to the lack of scalable, standardized methods enabling species-level assessment.

Ganglioside analysis is extremely challenging since (1) the glycan and lipid part exhibit highly converse chemical properties, (2) targeted extraction protocols are needed, (3) only a few standards are available, and (4) suitable glycolipid databases are still absent. Gangliosides exhibit amphiphilic properties since they consist of a sugar head group linked to a lipid subunit. Thus, they are neither fully covered by common glycomics nor lipidomics analytical workflows. Specialized glycolipidomics analytical workflows are required to bridge the gap between glycomics and lipidomics and decipher additional structural information. These glycolipidomics strategies have to deal with the complexity of two extremely heterogeneous classes: glycans and lipids. The theoretical number of glycan and non-saccharide permutations reaches almost Avogadro’s number(Laine, 1994; Merrill and Vu, 2016), raising the group of glycolipids among the most complex biomolecules from a combinatorial perspective. State-of-the-art approaches - including classical immunological, biochemical or thin-layer chromatography methods – entirely rely on class-specific ganglioside and glycolipid detection(Sarbu and Zamfir, 2018). For species characterization, they lack sensitivity or fail to provide detailed structural information, including the saccharide core (sialylation degree, anomers, branching) and the ceramide backbone (fatty acid, long-chain base, double bond position, and hydroxylation degree). Along the emergence of mass spectrometry-driven lipidomics, tailored ganglioside LC-MS workflows were proposed offering the possibility to collect both lipid and glycan information(Hájek et al., 2017; Ica et al., 2020; Sarbu and Zamfir, 2018). Predominantly, reversed-phase chromatography was reported as the method of choice(Ica et al., 2020; Sarbu and Zamfir, 2018), but also hydrophilic interaction chromatography was used for ganglioside class separation^30^. More than a decade ago, a seminal review pointed out three methodological milestones required to open the way for the field of glyco(sphingo)lipidomics: (1) automated online interfacing such as LC or CE with ESI-MS, (2) Ion Trap applications with MS^n^ capabilities, and (3) computational bioinformatics offering automated species annotation of spectra(Levery, 2005). While enormous advances were made regarding online class-specific separations prior to the mass spectrometric dimension, involving both high throughput UPLC and ion mobility, the latter two did not reach a satisfactory level yet. Despite the increased availability of sophisticated analytical workflows, the gangliosidome and glyco(sphingo)lipidome analysis remains a challenging task, typically conducted only by a few trained experts due to the tedious and complex manual annotation processes of fragmentation spectra.

In this work, we implemented a comprehensive and sensitive large-scale profiling strategy for gangliosides for the first time. The workflow is based on an online reversed-phase high-resolution MS^n^ ganglioside assay to analyze intact glycosphingolipids and annotate them up to the level of functional groups(Rampler et al., 2021). Using this novel LC-MS^n^ analytical strategy, all prevalent ganglioside classes are covered. Species annotation is based on platform-independent decision rules(Hartler et al., 2017, 2020). By applying this strategy to the analysis of MSCs induced towards differentiation in chondrogenic, adipogenic, and osteogenic lineages, we revealed the highest number of gangliosides reported in a single study so far(Hájek et al., 2017; Ica et al., 2020; Sarbu et al., 2016). Moreover, we demonstrated the capability of gangliosides as cell differentiation markers for mesenchymal stem cells.

## Results

### Surface marker expression and differentiation of human MSCs

MSCs have outstanding inherent therapeutically relevant properties, such as high in vitro proliferation capacities, secretion of biologically active components, and differentiation potential. The traditional paradigm of replacing damaged tissue by MSCs differentiation is currently challenged as MSCs were found to exert significant immunomodulatory effects on the innate and adaptive immune(Müller et al., 2021) system. For example, MSCs were successfully used to treat one of the most severe complications of COVID-19, which is the hyperactivation of the immune system, also known as cytokine release syndrome or cytokine storm, which can result in multiorgan failure and death(Song et al., 2021).

Until now, MSCs are characterized by the ability to differentiate towards the adipogenic, chondrogenic, and osteogenic lineage and the expression of several surface markers(Dominici et al., 2006). However, additional markers for specific cell states are needed for enhancing the quality control of cell-based therapy products.

Therefore, we characterized the ganglioside profile of native MSCs and MSCs differentiated towards the chondrogenic, adipogenic, and osteogenic lineage to find cell state-specific markers on a molecular species level. To confirm the identity of the adipose tissue-derived cells, the expression of several surface markers was assessed via antibody staining and flow cytometry. Conform to the minimal criteria of MSCs, we found CD73, CD90, and CD105 to be expressed, and CD14, CD20, CD34, and CD45 to be absent (**Figure 1a**). The MSCs were differentiated towards the chondrogenic, adipogenic, and osteogenic lineage using suitable media conditions. Histological staining confirmed the secretion of glycosaminoglycans in the chondrogenic samples, the formation of lipid vacuoles in the adipogenic samples, and the presence of calcium in the extracellular matrix of osteogenic samples that were cultured 21 days in the respective differentiation media (**Figure 1b**).

**Figure 1:**
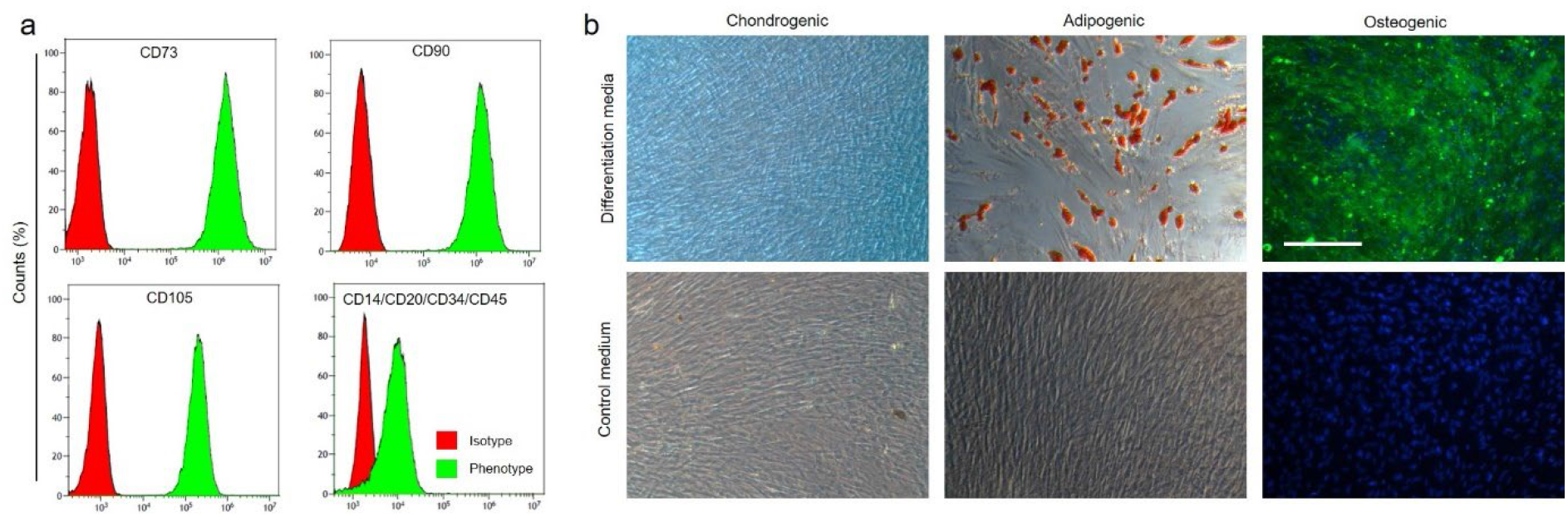
Characterization of human adipose-derived mesenchymal stem cells. **a** Flow cytometry histograms of adipose-derived MSCs stained for the respective surface markers after isolation in passage 2. **b** Micrographs of MSCs cultured in chondrogenic, adipogenic, osteogenic medium for 21 days and stained for glycosaminoglycans by alcian blue (chondrogenic, blue), lipid vacuoles by Oil Red O (adipogenic, red), or calcium deposition by calcein (osteogenic, green; counterstain with DAPI for cell nuclei, blue), respectively. Standard culture medium without differentiation supplements served as control. The scale bar represents 250 μm.

### Development of an automated RP-HRMS^n^ ganglioside assay

A novel ganglioside assay workflow was developed, enabling in-depth structural analysis and automated annotation of gangliosides (**Figure 2**). We bridge lipid and glycan analysis workflows by combining (1) an extraction protocol on methyl tert-butyl ether(Matyash et al., 2008), (2) reversed-phase high-resolution mass spectrometry and multi-stage fragmentation (RP-HRMS^n^), taking advantage of two parallel mass analyzers (Orbitrap, Ion Trap), and (3) a newly developed automated annotation workflow based on the open-source software Lipid Data Analyzer (LDA). Gangliosides were extracted from undifferentiated MSCs, and differentiated adipogenic, chondrogenic, and osteogenic cells. Concentrations were determined by protein content normalization and the addition of two deuterated internal standards (d5 GM3 36:1;O2, d5 GM1 36:1;O2). Reversed-phase ultra-high performance liquid chromatography was performed to separate the different ganglioside species based on their hydrophobic interaction with a C18 stationary phase. A standard acetonitrile isopropanol gradient was used to elute the gangliosides from the column, followed by lipid ionization in the heated electrospray source of the high-resolution mass spectrometer. In state-of-the-art HRMS workflows, glycosidic fragments are observed in negative mode only, whereas the ceramide part is monitored in positive mode only. In our workflow, the two mass analyzers (Orbitrap, Ion Trap) are used in parallel for multistage fragmentation (**Figure 2b**).

**Figure 2:**
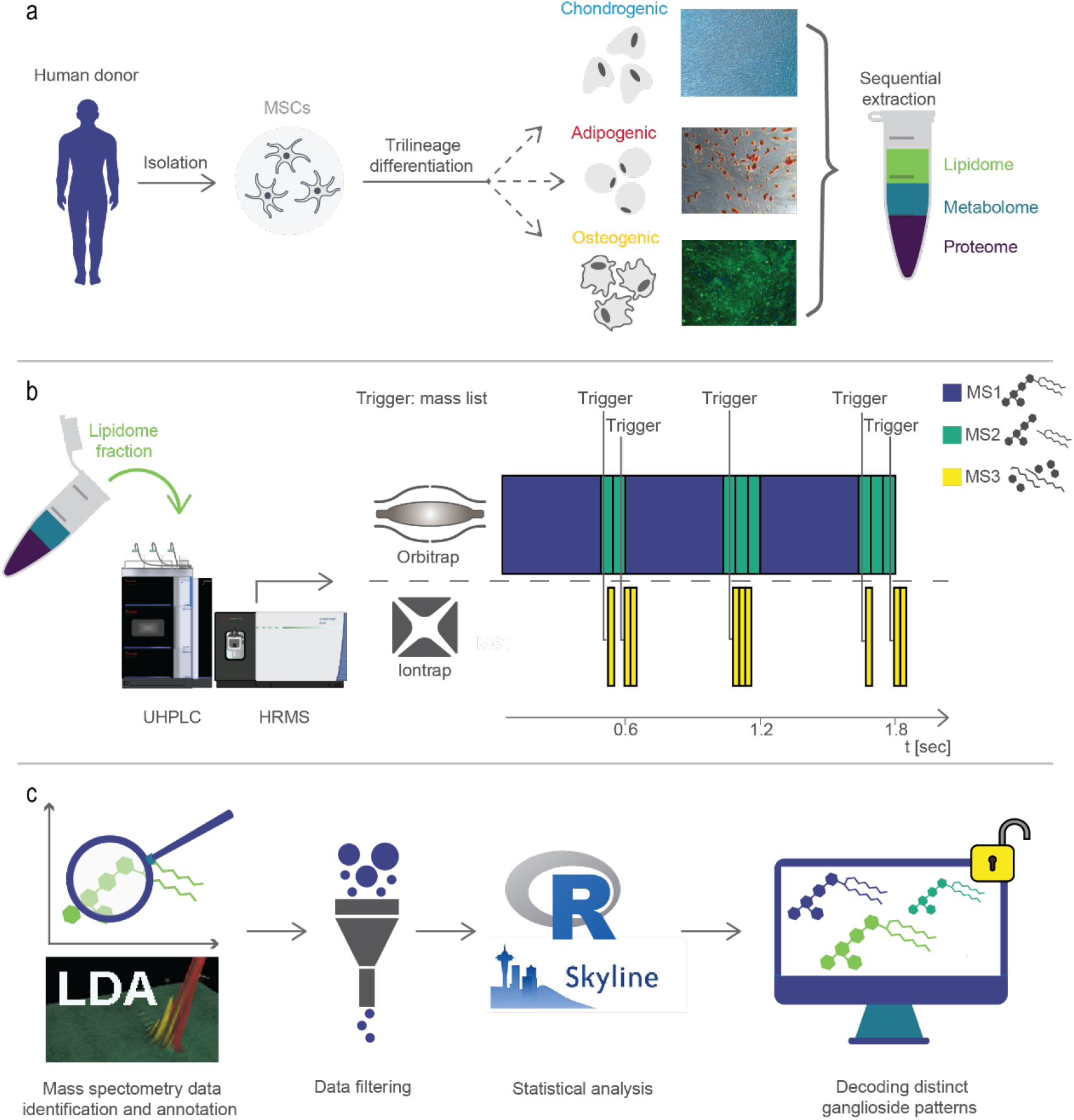
Developed glycolipidomics workflow to analyze different cell states. **a** Human MSCs were collected from surgical procedures and differentiated towards the adipogenic, chondrogenic, and osteogenic lineage. The lipid fraction was extracted, and **b** analyzed via RP-HRMS^n^ using multistage fragmentation with two parallelizable mass analyzers for in-depth characterization of the gangliosides. **c** Data analysis was performed using an open access workflow to decode distinct ganglioside patterns in native or differentiated MSCs.

This setup allowed for the acquisition of a set of fragments providing structural details for both glycosidic and lipid moieties in positive as well as in negative ionization mode (see **Figure 3**). The combination of MS2 and MS3 fragmentation permitted the complete structural characterization up to the lipid (molecular species) level(Liebisch et al., 2013, 2020) of all major ganglioside classes present in native MSCs and the three different cell lineages. The parallelized use of the mass analyzers enabled us to optimize the acquisition time towards higher information content (**Figure 2**).

**Figure 3:**
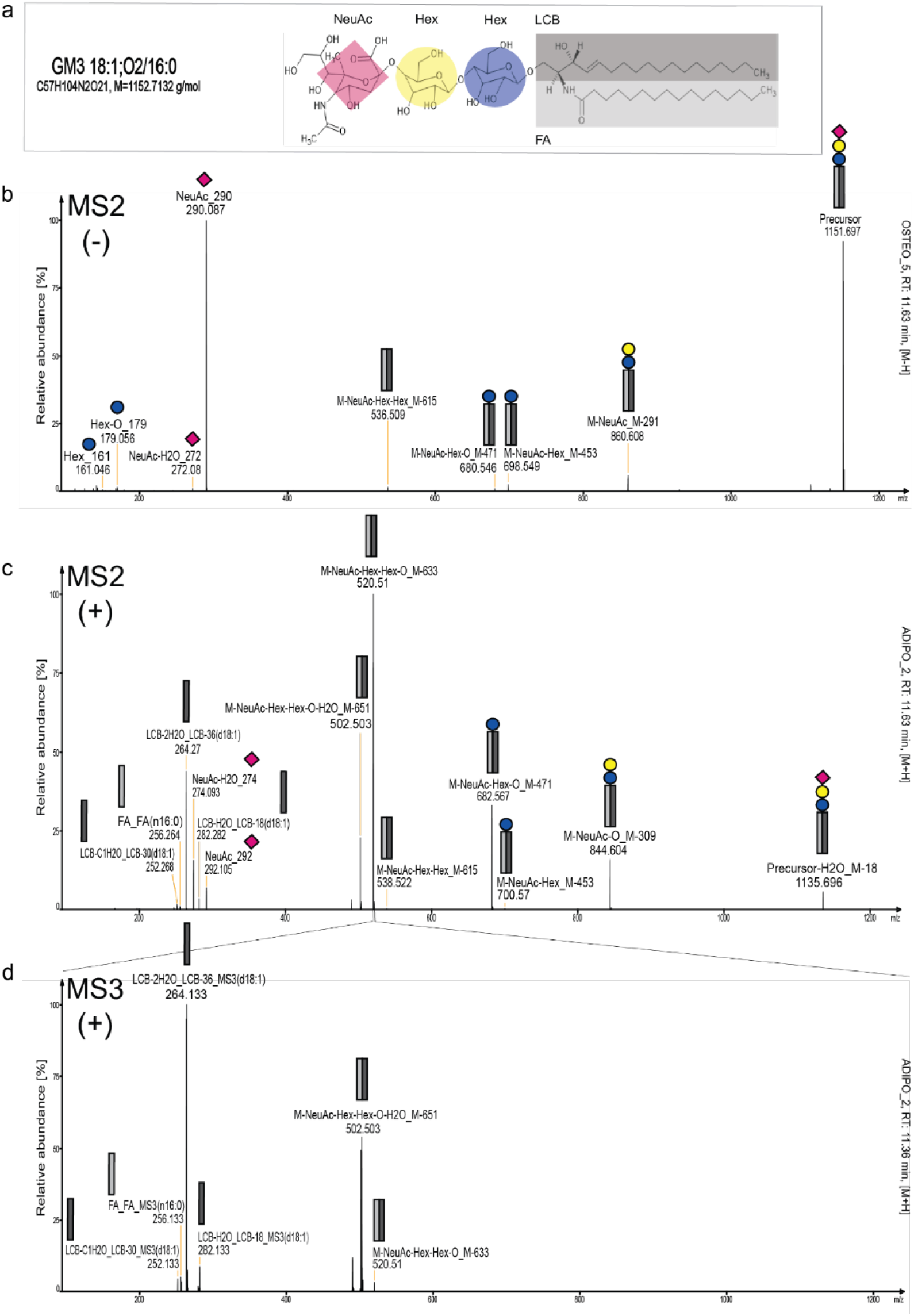
Molecular species ganglioside annotation by high-qualitiy spectra produced by the presented workflow. The annotation is based on fragmentation rules applied to the LDA software. http://genome.tugraz.at/lda2_review/lda_download.shtml **a** Gangliosides have a lipid moiety containing a long-chain base (LCB) and fatty acid (FA), shown as grey boxes. The glycan moiety (represented as blue and yellow circles) can vary in the number of hexoses (galactose and glucose; can not be distinguished by fragmentation spectra) and in the number of sialic acids (NeuAc, pink diamond). Spectral data of GM3 18:1;O2/16:0 in **b** negative (MS2) and positive (**c** MS2 and **d** MS3) ionization mode, respectively. In negative ionization mode, the fragments of the sugar part can be seen (Hex, NeuAc), as well as the intact precursor. In the positive ion mode, the composition of the ceramide part can be resolved.

Automated ganglioside annotation was performed based on in-house developed decision rule sets for the freely available software LDA optimized for sphingolipids LDA http://genome.tugraz.at/lda2.

To ensure data quality, we used additional fragment information corresponding to subunits specific for the different ganglioside classes. More specifically, we determined one mandatory MS2 fragment followed by MS^n^ fragment assignment based on rules for the glycan (e.g., number of sialic acids) and lipid part (LCB and FA moiety, hydroxylation state) for all ganglioside classes detected. A simplified overview of the final glyoclipidomics workflows can be found in **Figure 2**. Using the presented strategy, we were able to identify 263 unique gangliosides (on species-level e.g. GM3 36:1;O2) of the classes (**Supplementary Figure 1**) GM1, GM2, GM3, GM4, GD1, GD2, GD3, GT1, GT2, GT3, GQ1, and GP1 (**Supplementary Table 2**) in MSCs and differentiated cell line samples. Out of the 263 annotated gangliosides, 224 species from eight classes (GM1, GM2, GM3, GM4, GD1, GD2, GD3, GT3) were quantified down to the fmol level, based on one-point calibration with the deuterated internal standards providing concentration estimations in nmol per mg protein. The annotation was confirmed using ganglioside standards (total ganglioside mixtures, ganglioside class, and deuterated single standards) and corresponded to category D quality (meaning species or molecular species level were determined)(Rampler et al., 2021). Using our novel workflow, fully automated lipid species annotation was possible, which is a significant advance to state-of-the-art ganglioside analysis workflows. The most abundant gangliosides determined were in the nmol per mg protein range in all samples, where GM3 was the class of highest abundance, followed by GM2 and GD3, respectively, which are classes containing exactly one sialic acid (**Supplementary Table 1**, **Supplementary Figure 2**).

### Ganglioside-specific cell state differences and their potential as differentiation markers

In the context of therapeutic applications, it is crucial to maintain the cellular identity and functionalities during *ex vivo* culture to ensure the safety and efficacy of MSCs. However, extensive *ex vivo* expansion of cells is necessary to obtain sufficient cell numbers for therapeutic treatments. Due to frequent passaging on traditional polystyrene surfaces, such as Petri dishes, T-flasks, or well-plates, genetic alterations accumulate over time, resulting in the loss of relevant therapeutic properties(Capelli et al., 2014; Hladik et al., 2019; Miura et al., 2006). This can impair the therapeutic outcome and pose a safety issue as malignant transformations can be expected(Wang et al., 2013). Therefore, additional markers are desirable to monitor and define the stem cell phenotype during *ex vivo* culture. In our previous work, the adipogenic differentiation of MSCs, metabolite, and lipid non-targeted data analysis revealed gangliosides as significantly regulated compounds for MSCs compared to adipocytes(Rampler et al., 2019). Here, we differentiated MSCs towards the chondrogenic and osteogenic lineage to investigate the general ganglioside marker potential. Using our optimized RP-HRMS^n^ assay followed by automated ganglioside annotation, we detected the intact glycosphingolipids and deduced both glycan and lipid structural information. Classical immunological, biochemical, or thin-layer chromatography methods rely on class-specific ganglioside detection. All lipid backbone variations are summarized in a specific glycan-dependent ganglioside class (e.g., GM3, GM4, etc.). **Figure 4** shows the total ganglioside concentration in the different cell lineages, e.g., mimicking the low information content provided by a state-of-the-art ganglioside class-specific antibody assay. Although specific trends can be observed, such as (1) higher glycan series in osteocytes or (2) a significantly lower expression of GM2 in MSCs compared to the differentiated cells, important structural information is unavailable. It is not possible to single out the relevant ganglioside species and derive more details about their specific biological functions to enhance the quality control of MSC-based therapy products.

**Figure 4:**
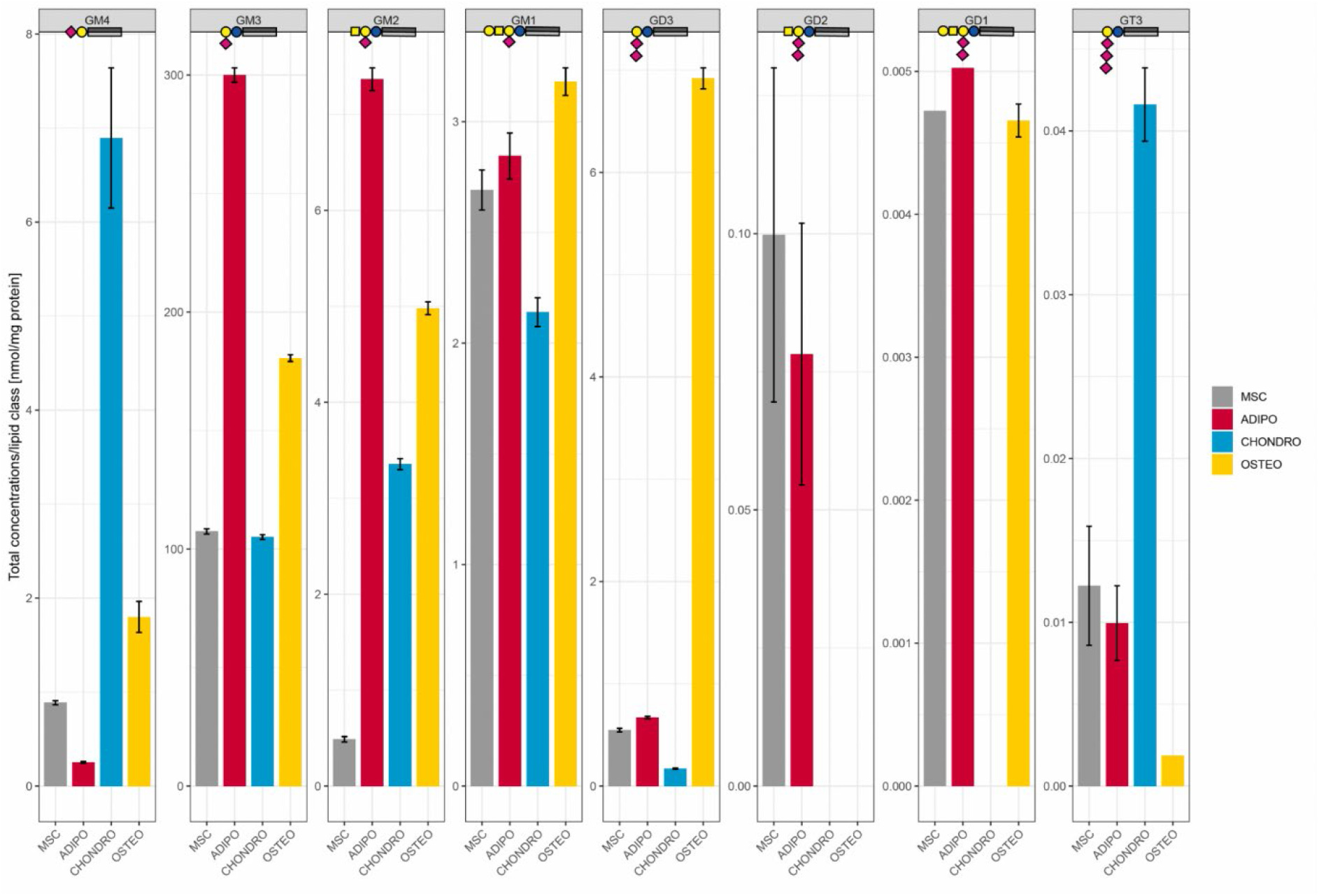
Lipid class overview according to detected sialic acid length in MSC and differentiated cell lineages. We can observe an upregulation trend of GD3 in osteocytes. Considering the concentration on the class level only, GM3 is highest in adipocytes. However, at the fine-grained lipid species level, we detected cell type-specific differentiation patterns between all monitored tissue types. A distinct effect of increasing sialic acid length could not be observed, further highlighting that the discriminative power is hidden in the specific ceramide moieties. The shown class concentrations were derived by the sum of intensities of the individually identified lipid species ([M-H] adduct).

In this study, we enhanced the level of ganglioside annotation up to the molecular species level by exploiting the benefits of multistage fragmentation. The 263 detected gangliosides in MSCs, adipogenic, osteogenic, and chondrogenic differentiated cells contained ceramide moieties ranging from 30:0 to 48:2 including both dihydroxylated and trihydroxylated species, and the highest abundance showed 34:0;O2; 34:1:O2, 40:1;O2, and 42:1;O2 (**Supplementary Table S2**). The highest abundance long-chain base was expectedly sphingosine (18:1:O2) in all cell lineages. Most LCBs were dihydroxylated with a length between 16-20 carbons, with a predominance for even carbon numbers. If dihydroxylated species were observed, chances of observing the corresponding lower abundant trihydroxylated LCB (e.g. 18:1;O3, 18:2;O3) were higher.

Successful isomer separation was possible by hydrophobic interaction with the lipid part on the reversed-phase column followed by potential molecular species assignment using the MS^n^ approach as shown by the example of GM3 18:1;O2/24:1 (retention time: 19.42 min) - upregulated in adipocytes - and its isomer GM3 18:2;O2/24:0 (retention time: 19.98 min) upregulated in osteocytes (**Supplementary Figure 3).** Interestingly, the total number of ganglioside annotations increased from mesenchymal stem cells (50 gangliosides) to all differentiated states (Adipocytes: 70, Chondrocytes 130, Osteocytes: 160) (**Supplementary Figure 4**). Although we could observe some differences in lipid content of other lipid classes -e.g., TG, PC, and PE were upregulated in adipocytes, compared to the other cell lineages (**Supplementary Figure 5**)- the overall molecular lipid species remained constant in the bulk lipid classes. In contrast, the gangliosides patterns revealed increasing annotation numbers upon differentiation and enabled successful cell lineage separation based on distinct ganglioside species (**Figure 5 (b)**). Out of the 263 detected ganglioside species, 53 belonging to the classes GM3, GM1, GM2, GD3 exhibited significant differences (p-value< 0.05, Tukey’s HSD, Metaboanalyst(Chong et al., 2018)) in the various cell lineages (**Figure 5**). Using these 53 candidate markers, the distinction between human MSCs, adipocyte, osteocyte, and chondrocyte sample groups (n=5) was easily possible (**Figures 5+6**). For the ganglioside marker selection, we annotated the molecular species level based on MS2 and MS3 fragmentation information whenever possible (dependending on ganglioside concentration and quality of spectra). The increased separation and assignment potential of the presented RP-HRMS^n^ approach revealed 62 lipid species hits (**Supplementary Figure 2**), including isomeric molecular lipid species markers specific for particular cell states (e.g., **Supplementary Figure 3**).

**Figure 5:**
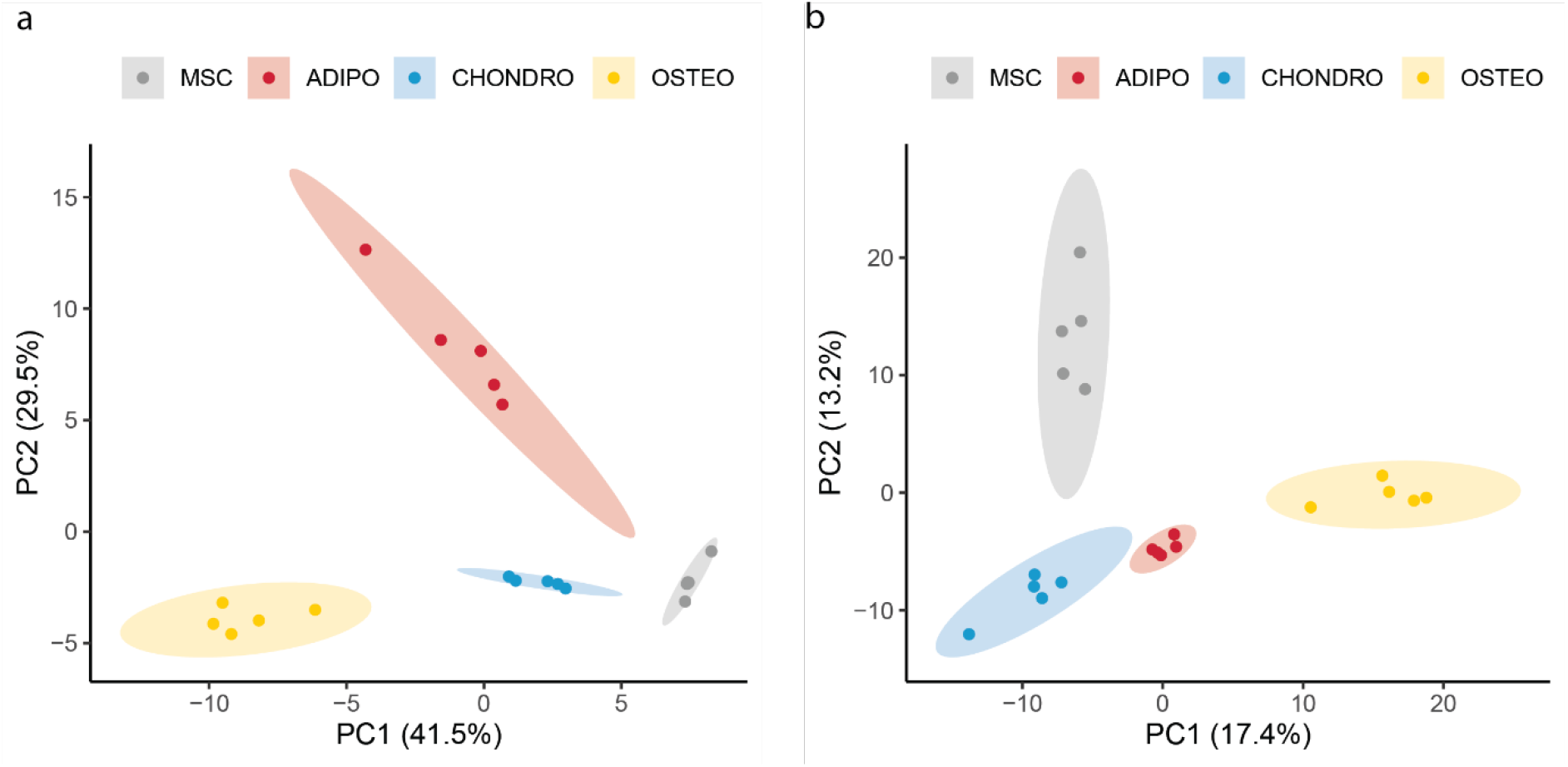
Principal component analysis (PCA) of identified gangliosides separates MSC and differentiated cell lineages. **a** PCA indicates that 53 unique marker candidates (lipid species level) explain 71% of the total variance in PC1 and PC2 and allow a clear distinction of MSC and each of their differentiated cell lineages (n=5); this corroborates the high potential of the selected candidates to serve as differentiation markers. **b** PCA of all 263 identified gangliosides also allows separation of cell lineages, with 30.6% of the variance explained by PC1 and PC2.

**Figure 6:**
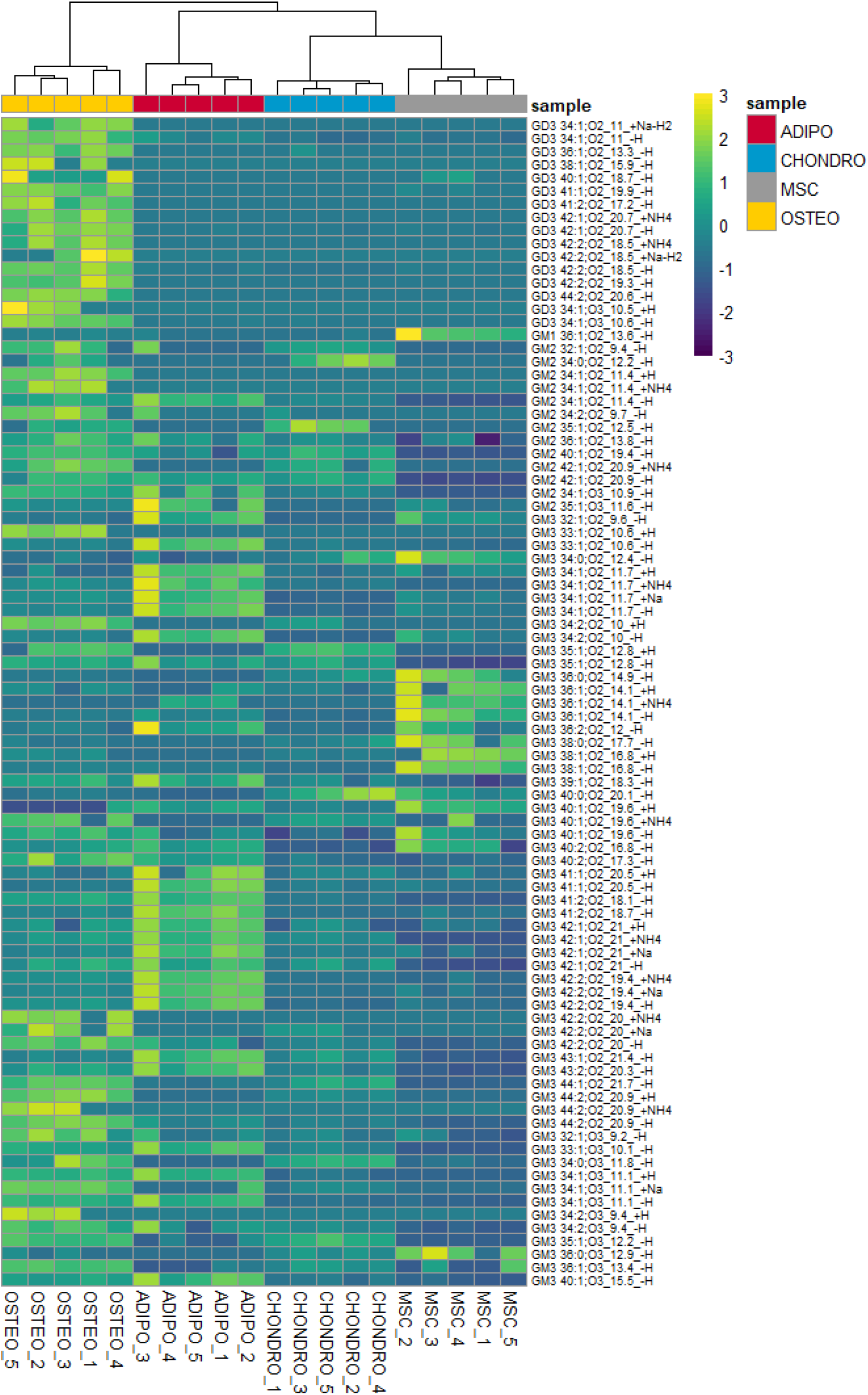
Heatmap of potential marker candidates. Biological sample replicates(n=5) cluster according ganglioside regulation patterns observed in both polarities. Sample names of the different gangliosides include the lipid species, the retention time (in minutes), and the adduct. Hierarchical clustering was performed using correlation distance (euclidean) andaverage linkagescale (ward.D2). The heatmap was plotted using the R packagepheatmap.

Confirming our previous data on adipogenesis(Rampler et al., 2019), GM3 18:1;O2/16:0 was upregulated only in adipocytes. Generally, the class of GM3 contained 21 signficantly regulated adipocyte markers with nine GM3 species (GM3 18:1;O2/14:0, GM3 17:1;O2/16:0, GM3 18:1;O2/16:0, GM3 18:2;O2/16:0, GM3 18:1;O2/23:0, coeluting GM3 18:1;O2/23:1 & GM3 17:1;O2/24:1, GM3 18:1;O2/24:0, GM3 18:1;O2/24:1, colueting GM3 18:1;O2/25:0 & GM3 19:1;O2/24:0 & GM3 20:2;O2/23:0) highly upregulated compared to all other cell lines. Despite the observation that the class of GM3 is generally upregulated in adipocytes (**Figure 4**), GM3 also contained gangliosides - GM3 36:0;O2, GM3 18:1;O2/18:0, GM3 16:0;O2/22:0, GM3 18:1;O2/20:0, GM3 36:0;O3 - highly upregulated in MSCs suggesting these lipids as potential non-differentiation/stemness markers (**Figure 7**). Additionally, one GM1 (GM1 18:1;O2/18:0) was also upregulated in MSCs compared to the other cell lines (**Figure 7**). Conversely, GM2 ganglioside species were generally downregulated or completely absent in MSCs compared to all other differentiated cell lines, indicating GM2s as differentiation marker (**Figure 4+7**). In chondrocytes, we observed a general increase of saturated gangliosides, which was significant compared to all other sample groups, specifically in GM2 34:0;O2 and GM3 40:0;O2. Additionally, GM2 35:1;O2 was upregulated, whereas GM3 16:1;O2/24:1 (at RT 16.8 min) was downregulated in chondrocytes compared to its chromatographically separated isomer GM3 18:2;O2/22:0 (at RT 17.3 min) which is upregulated in osteocytes (**Figure 7**). In osteocytes, a general increase of all significantly regulated GD3 gangliosides (GD3 18:1;O2/16:0, GD3 18:1;O2/18:0, GD3 18:1;O2/18:0, GD3 40:1;O2, GD3 41:1;O2, GD3 41:2;O2, GD3 18:1;O2/24:0, GD3 18:1;O2/24:1, GD3 42:2;O2, GD3 44:2;O2, GD3 18:1;O2/16:0) was observed (**Figure 7**). A general trend towards longer glycan and ceramide parts in the class of GM3 and GD3 as well as trihydroxylated GM3s (GM3 32:1;O3, GM3 34:0;O3, GM3 18:2;O3/16:0, GM3 35:1;O3, GM3 36:1;O3) is obvious in osteocytes as shown in both the class-specific detection trend in **Figure 4** and the marker candidate list depicted in **Figure 7**.

**Figure 7:**
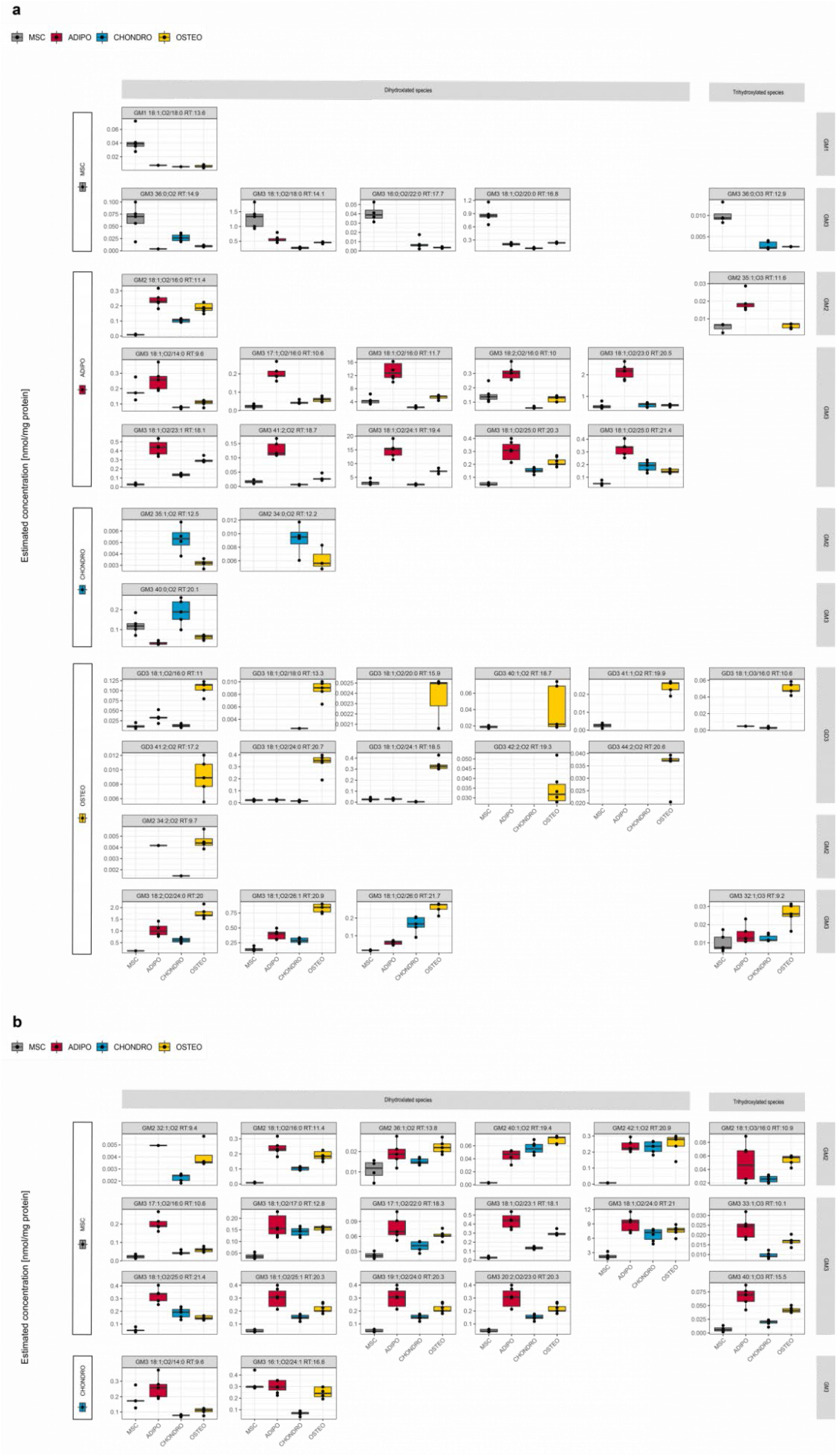
Box plots of significantly regulated potential ganglioside differentiation marker for each cell type. Marker candidates are separated into di- and trihydroxylated ganglioside species and the corresponding ganglioside class. Estimated concentration are based on internal standardization with GM3 d5 18:1;O2/18:0 and normalization to the protein content ([M-H] adduct). **a** Upregulated gangliosides for MSC and each differentiated cell lineage. **b** Downregulated gangliosides for each MSC and each differentiated cell lineage.

Out of the 53 individual ganglioside markers, 38 were successfully resolved on the molecular lipid species level using MS2 and MS3 fragment information. From this panel, 16 novel gangliosides (GM3 17:1;O2/16:0, GM3 18:2;O2/16:0, GM3 18:2;O2/18:0, GM3 16:0;O2/22:0, GM3 17:1;O2/22:0, GM3 16:1;O2/24:1, GM3 18:2;O2/22:0, GM3 17:1;O2/24:1, GM3 18:2;O2/24:0, GM3 19:1;O2/24:0, GM3 19:1;O2/24:1, GM3 20:2;O2/23:0, GM3 18:2;O3/16:0, GM3 18:1;O3/16:0, GD3 18:1;O3/16:0, GM2 18:1;O3/16:0) were reported on the molecular species level (marked in blue in **Supplementary Table 1**). We defined a lipid molecular species as ‘novel’ if neither present in ChEBI(Hastings et al., 2016), HMDB(Wishart et al., 2009), LIPID MAPS(Sud et al., 2007), SwissLipids(Aimo et al., 2015) or the recent comprehensive ganglioside lists published in the literature.(Ica et al., 2020; Sarbu et al., 2016) The fact that from the subset of 38 molecular lipid species almost half of the gangliosides (16) were never reported so far proves the general power of the presented RP-HRMS^n^ approach to identify unknown gangliosides in samples of interest with unprecedented structural detail.

## Discussion

In this work, we established a novel ganglioside assay based on reversed-phase high-resolution mass spectrometry and multi-stage fragmentation. Essential aspects of the presented assay are sensitivity down to fmol ranges complemented by an automated open-source annotation workflow by LDA, which was recently tailored to sphingolipid detection(Hartler et al., 2020). By (1) applying decision rule sets from both MS2 and MS3 spectra deduced from our experimental data in standards and samples, (2) making use of positive and negative ionization mode data, and (3) including retention time information, we significantly improved the ganglioside annotation coverage and quality.

We applied our novel glycolipidomics assay to characterize the ganglioside profile of native MSCs and MSCs differentiated towards the chondrogenic, adipogenic, and osteogenic lineage in order to find cell state-specific markers on a molecular species level. Here, we reported the highest number of gangliosides so far, encompassing 263 unique lipid species. Cell state-specific markers for native or differentiated MSCs could be used as additional markers for quality control of cell-based therapy products. The cells in this study fulfilled the minimal criteria for MSCs, including the positive expression of CD75, CD90, and CD105 and the negative expression of CD14, CD20, CD34, and CD45. Furthermore, the cells were successfully differentiated towards the chondrogenic, adipogenic, and osteogenic lineage. This study’s findings are in accordance with observed ganglioside regulations of our previous study on MSC and adipogenic cell differentiation coming from a different stem cell donor source (female, other adipose tissue part)(Schoeny et al., 2021).

As gangliosides are present on the membrane’s outer leaflet and influence the membrane curvature and segregation(Sonnino et al., 2007), they represent accessible surface markers that influence cell morphology, being necessary for both stem cell stiffness and cell differentiation. Thus, gangliosides may represent excellent candidates to serve as stem cell-state markers(Bergante et al., 2014). However, a structured analysis of the ganglioside profile of native and differentiated MSCs is still missing. Other studies investigated the presence or effect of gangliosides on the level of ganglioside classes GD2(Bergante et al., 2014; Freund et al., 2010; Martinez et al., 2007) but not at the detailed ganglioside species/molecular species level. With our sensitive profiling approach, we were able to identify many promising potential biomarker candidates with increased structural information. Nonetheless, some of the detected candidate species may be attributed to the donor variability (age, sex, and environmental factors), which affect relevant MSC properties(Heathman et al., 2016; Mushahary et al., 2018). Furthermore, the cell culture conditions (media and media supplements, culture format, 2D/3D, oxygen level, etc.) may have detrimental effects on all levels of cellular behavior and expression patterns. While these confounding variables do not reduce the analytical power of our assay, to confirm whether our candidates are generally applicable markers for MSC differentiation, we are aware that a comprehensive analysis of the ganglioside profile during differentiation of MSCs from different tissues and donors of different ages and genders is necessary. Those limitations mentioned above are potential causes for differences to previous studies, e.g., former studies reported differential expression of GM1, GM3, GD1, and GD2(Bergante et al., 2018; Lee et al., 2010; Martinez et al., 2007) in MSCs from different sources. While our results confirm the presence of these classes in undifferentiated MSCs, we found that GM1 is similarly abundant during adipogenic, chondrogenic, and osteogenic differentiation, and GM3 is higher expressed in adipogenic than in osteogenic differentiated MSCs. GD2 is not expressed in chondrogenic or osteogenic cells but in MSCs and adipogenic cells (**Figure 4**). Thus, analysis on the level of ganglioside classes did not result in a single class that could serve as a good marker for stemness. However, we identified several GM3s (e.g., GM3 18:1;O2/18:0, GM3 16:0;O2/22:0, GM3 18:1;O2/20:0, GM3 36:0;O3 18:1;O2/18:0, GM3 18:1;O2/20:0) as promising markers for native MSCs. Regarding the adipogenic differentiation of MSCs, we confirmed our prior finding of an increase of GM3, which is known to be expressed in terminally differentiated adipocytes from adipose tissue(Nagafuku et al., 2015) and is recognized to participate in insulin signaling. The accordance of the GM3 increase is of special interest as different MSC donors and materials were used in this work and our previous study(Nagafuku et al., 2015). Hence, we propose several GM3 ganglioside (molecular) species as potential markers for adipogenic differentiation (**Figure 7**). Regarding chondrogenic differentiation, our study does not confirm former results(Ryu et al., 2020). Instead of increased expression of GM3 and GD3, we found ganglioside classes GM4 and GT3 significantly upregulated in the chondrogenic lineage compared to native MSCs. On the lipid species level molecular level, GM2 d34:0 and GM3 d40:0 were upregulated considerably in chondrogenic differentiated cells, compared to native MSCs and the differentiated cells. In the osteogenic differentiation of MSCs, we can confirm a significant increase in GM1 expression (**Figure 4**) as observed before(Bergante et al., 2018). Interestingly, we also found a massive increase in the class of GD3 in osteogenic cells, including several potential GD3 markers which could be characterized on the molecular species level (**Figure 4**). Overall, some of the literature observations match, whereas others are not confirmed. This can be explained due to MSCs from different sources, age, sex, and culture conditions, as well as the greater structural details we can provide using our RP-HRMS^n^ ganglioside assay. While the GD3 upregulation on the class level is also observed in differentiated osteogenic cells in literature(Moussavou et al., 2013), most of our examples reveal candidate marker on the (molecular) lipid species level as shown for the class of GM3 with distinct ganglioside markers for native MSCs, and MSCs differentiated towards the adipogenic, osteogenic and chondrogenic differentiation, respectively. As state-of-the-art biochemical and immunological analysis methods only monitor the ganglioside class level, they are not sufficient to resolve the complex ganglioside pattern.

This work clearly demonstrated that ganglioside profiles could be autonomously used as surrogate markers throughout the whole differentiation states of the three chosen lineages. In-depth structural characterization using the RP-HRMS^n^ assay was possible by providing both ceramide and glycan information. Therefore, monitoring the ganglioside profile might be used as a quality control parameter for MSC differentiation in tissue engineering processes and for maintaining stemness during MSC expansion to manufacture cell therapy products. For example, CD markers are defined cell surface molecules that act as targets for immunophenotyping to determine the cell state. Interestingly, the sugar moiety of GD3 serves as such a target for quality control and is defined as CD60(Zhang et al., 2019). In the context of the presented results, we propose to include molecular-level ganglioside-specific markers or marker sets to characterize stem cell differentiation processes. Our results suggest gangliosides as potent markers for human stem cell differentiation into various cell lineages confirming the previous findings(Bergante et al., 2014; Yanagisawa, 2011). Although gangliosides can influence membrane shape(Sonnino et al., 2007) and show differentially expressed patterns in different cell types(Freund et al., 2010), little is known to what extent ganglioside patterns affect and/or trigger stemness and stem cell differentiation. Monitoring both the sugar and lipid moiety of gangliosides paves the way for detailed investigations of these important interactions. Sophisticated glycolipidomics approaches enable in-depth molecular characterization of gangliosides and glyco(sphingo)lipids. High-resolution MS-based glycan-library screening was recently performed to determine glycolipid binding upon viral entry of SARS-CoV-2 and proved an interaction preference for monosialylated gangliosides(Nguyen et al., 2022). In this workflow, the glycan part was released to determine the sugar part of the interacting gangliosides. Using our presented LC-MSn assay, direct read-out of native gangliosides is possible without removing the glycan part. The multidimensional separation based on chromatography and multistage fragmentation enables both (1) ganglioside identification of classes and isomeric differences as well as (2) quantitative comparison between sample groups. Our proposed workflow can be seen as the hallmark for shedding light on the enigmatic processes of gangliosides and glycolipids in general, which are critical mediators in cell-cell interaction and signaling pathways. Hence, we believe multidimensional separation and HRMS^n^ based assays will lay the foundation to unravel how glycolipids orchestrate the fatty sweet symphony in cells and organisms. Our workflow is an innovative strategy to determine and make use of novel glycolipid-based CD markers for stem cell phenotype characterization.

## Methods

### Cell culture

The use of human tissue was approved by the ethics committee of the University of Lübeck (EK Nr: 20-333), and the donor (male, 29 years) provided written consent. MSCs were isolated within approximately 12 h after surgery as previously described(Egger et al., 2017). MSCs were cultured in a standard medium composed of MEM alpha (Thermo Fisher Scientific, Waltham, MA, USA), 0.5% gentamycin (Lonza, Basel, Switzerland), 2.5% human platelet lysate, and 1 IU/ml heparin (both PL BioScience, Aachen, Germany) on standard T-flasks (Sarstedt, Nümbrecht, Germany), in a humidified incubator at 37°C and 5% CO2. After confluency was reached, cells were detached with accutase (Sigma Aldrich) and cryo-preserved in liquid nitrogen as previously described(Neumann et al., 2014). Upon use, MSCs were thawed and subcultivated in T-flasks once, resulting in passage two.

### Differentiation

For MSC differentiation, 4,000 cells/cm^2^ in passage two were seeded on a 6-well plate (Sarstedt) coated with fibronectin (2 μg/cm^2^; Sigma Aldrich) and allowed to grow confluent. The medium was changed to adipogenic, chondrogenic, or osteogenic medium (Miltenyi Biotech, Bergisch Gladbach, Germany, supplemented with 0.5% gentamycin, n=6 replicates for each medium). Cells were cultivated for 21 days, and the medium was changed every three to four days. The confluent MSCs at day zero of differentiation served as control.

### Histological stainings

MSCs from adipogenic and chondrogenic mediums were fixed with 4% paraformaldehyde (PFA, both Sigma). Cells cultured in adipogenic differentiation medium were stained with Oil Red O (Sigma Aldrich) for lipid vacuoles. Cells were stained for glycosaminoglycans with Alcian blue (Sigma Aldrich) to confirm chondrogenic differentiation. MSCs cultivated in the osteogenic medium were fixed with 96% ethanol and stained for calcium with Calcein and nuclei counterstained with DAPI (Sigma). Cells at day zero served as control.

### Phenotyping

To determine MSC surface marker expression, MSCs in passage two were detached by accutase treatment and stained with a human MSC phenotyping kit and an anti-HLA-DR antibody (both Miltenyi Biotec) according to the manufacturer’s instructions. In this kit, the antibodies for the negative markers (CD14, CD20, CD34, and CD45) are labeled with the same fluorophore to generate a negative marker panel. According to the manufacturer’s manual, an approximately 10-fold increase of the fluorescence intensity of the negative markers is expected for negative samples compared to the isotype control. The stained cells were resuspended in a suitable volume of flow cytometry buffer (0.5% fetal bovine serum, two mM EDTA in PBS), and one × 10^4^ gated events per sample were recorded on a CytoFLEX S (Beckman Coulter, Brea, CA, USA). Subsequent analysis was performed with Kaluza Flow Cytometry software (version 1.3, Beckman Coulter).

### Glycolipidomics

#### Standards and solvents

All solvents were LC-MS grade. Ganglioside standards were from Cayman Chemical (GD2, GM4, Ann Abor, USA) or Avanti Polar Lipids, Inc. (GM3 bovine milk, total ganglioside extract from porcine brain, Alabaster, Alabama, USA) and were weighed and dissolved in an appropriate solvent (IPA/H2O (65%/35%, v/v). Deuterated GM3 18:0/18:1;O2-d5 and GM1 18:0/18:1;O2-d5 standards were purchased from Avanti Polar Lipids, Inc. (Alabaster, Alabama, USA) and used as an internal standard for further analysis.

#### Ganglioside extraction

An adapted SIMPLEX protocol(Coman et al., 2016) was used to extract gangliosides as described previously(Rampler et al., 2019) since it enables the simultaneous collection of lipids (upper phase), metabolites (lower phase), and the protein pellet. In short, MSC and differentiated cell lineages (adipocytes, chondrocytes, and osteocytes) were quenched directly on the 6-well plate before storage at −80 °C until sample preparation. Deuterated internal standards were added directly to the samples to compensate for losses during extraction. Harvesting of adherent cells (~2*10^5^ cells/well) was performed using a cell scraper. Subsequent extraction was accomplished using a mixture of cold methanol, methyl-tert-butyl ether (MTBE), and ten mM ammonium formate. Five replicates/conditions, as well as four medium blanks (control), were prepared. The lipid fractions were collected, dried, and reconstituted in IPA/H2O (65%/35%, v/v). To determine the protein concentration, the BCA assay (Pierce kit, Thermo Fisher) was applied.

#### Ganglioside profiling with RPLC-HRMS^n^

A tailored ganglioside method was developed using a Vanquish Horizon UHPLC system coupled via heated electrospray-ionization (HESI) to a high-field Orbitrap ID-X™ Tribrid™^1^ mass spectrometer (both from Thermo Fisher Scientific). RP chromatography was accomplished using an Acquity HSS T3 (2.1 mm × 150 mm, 1.8 μm, Waters) column with a VanGuard pre-column (2.1 mm × 5 mm, 100 Å, 1.8 μm). The column temperature was set to 40 °C, the flow rate was set to 0.25 μL min^-1^, and the injection volume was set to 5 μL. The injector needle was flushed with 75% isopropanol (IPA) and 1% formic acid (FA) in between the injections. Acetonitrile (ACN)/H2O (3:2, v/v) was used as solvent A and IPA/ACN (9:1, v/v) as solvent B, both containing 0.1% formic acid and 10 mM ammonium formate. A gradient of 30 minutes under following conditions was applied: 0.0-2.0 min 30% B, 2.0-3.0 min ramp to 55% B, 3.0-17.0 min ramp to 67% B, 17.0-22.0 min ramp to 100 %, 22.0-26.0 min 100% B, 26.0 min fast switch to 30% B and 26.0-30.0 equilibration at starting conditions (30% B).

The ESI source parameters were as follows: 3.5 kV (positive ion mode) and 3.0 kV (negative ion mode), sheath gas 40, auxiliary gas 8, sweep gas 1, capillary temperature (ion transfer tube temperature) 275 °C, auxiliary gas heater (vaporizer temperature) 350 °C, radio frequency (RF) level 45%. Positive and negative ionization mode data were acquired in separate runs.

Spectral data were acquired in profile mode. For full MS runs, a mass range of *m/z* 500-2,000 at a resolution of 120,000 was selected. The automatic gain control (AGC) target was set to standard, the maximum injection time (MIT) was 100 ms. MS2 and MS3 scans were performed with data-dependent acquisition (DDA). For MS2 scanning, a top 5 method with a resolution of 15,000 was applied. The normalized collision energy (NCE) was set to 23 (+) and 27 (-) (HCD activation), respectively, the isolation window to *m/z* 1.5, the AGC target to standard, and the MIT to 60 ms. The dynamic exclusion of triggered *m/z* was set to 5 s, and a ganglioside-specific inclusion list was implemented. For MS3 spectra, the mass range was reduced to *m/z* 300-800 to gain fragments of the ceramide moieties (LCB, FA) to elucidate the molecular lipid species composition. The Ion Trap was selected as a mass analyzer with a fixed collision energy of 30 % (CID activation), with 10 ms activation time and an activation Q of 0.25. The Ion Trap scan rate was set to rapid, the isolation window to *m/z* 1.5 (MS1) and 2.0 (MS2), respectively, the AGC target was set to standard and MIT to automatic. The MS3 scans in the Ion trap were parallelizable with MS1 and MS2 scans in the Orbitrap, increasing information content and saving time (**Figure 2**).

Deep ganglioside and lipid profiling was performed on sequential automated exclusion lists (including blank subtraction) enabled by AcquireX data acquisition software within the Orbitrap ID-X™ Tribrid™ mass spectrometer.

### Data evaluation

The ganglioside assignment was performed using the LDA (version 2.8.2)(Hartler et al., 2011, 2020), the *OrbiTrap_IDX_heavy* settings were optimized and can be found in the provided LDA version LDA http://genome.tugraz.at/lda2 including necessary decision rule sets and corresponding mass lists. Analysis results were exported separately for positive and negative ion mode using the rdb export option. Areas of internal standards were received from Skyline (version 21.1) and exported as a csv file. LDA (ganglioside annotations), Skyline (internal standard), and BCA (protein content) results were then evaluated using R Studio (version 4.1.2). All annotation hits were normalized to the corresponding adduct of the internal standard for both polarities. The lipid species’ concentration was estimated via one-point calibration using the known concentration of the internal standards. Subsequently, normalization to the protein content was performed. Several quality control filters were applied, including (1) retention time and elution order using the equivalent carbon number (ECN) model(Holčapek et al., 2015), (2) area threshold, and (3) retention time matching of adducts from both polarities. Finally, after automated LDA annotation and filtering (applied filters: RT 2-26 min, at least one verified MS2 spectrum, area > 30 000), for the regulated markers, additional stringent quality criteria (a. MS1 peak shape, MS2 spectra; b. each trihydroxylated ganglioside has a corresponding dihydroxlyated species with fitting RT, c. ECN model, d. adducts) were applied manually.

Data was exported to MetaboAnalyst(Chong et al., 2018) (version 5.0) R Studio was used for statistical analysis (anova, heatmap, PCA). For significantly regulated species (potential marker candidates), the retention times were manually verified for plausibility (ECN model plus trihydroxylated species prior to dihydroxylated ones).

## Supporting information

Supplementary Table 1: Overview of gangliosides marker

Supplementary Table 2: Annotated gangliosides in native and differentiated MSCs.

## Acknowledgments

This work was supported by the University of Vienna, the Faculty of Chemistry, and the Vienna Metabolomics Center (VIME). We are grateful to PD Dr. med. Maike Keck for providing the MSC source tissue.

The authors thank all members of the Rampler, Koellensperger lab (University of Vienna), Hartler lab (University of Graz), and Kasper lab (BOKU Vienna) for a great team spirit and scientific exchange. We further acknowledge our technicians Christoph Baumgartinger, Petra Voljecnik, and Julia Zoller für continuous lab support. We also thank Ilias Nikolits for the help with the graphical workflow representation.

## Data availability

All data are included in the article and the **Supplementary Information** or are available from the authors upon request. The open-access version of LDA for ganglioside analysis will be freely available upon publication, as well as the (1) raw data, (2) mzml files, and (3) general ganglioside annotation list at GNPS (WinSCP client has to be downloaded, ftp://MSV000089064@massive.ucsd.edu).

## Ethics declaration

The isolation of MSCs from human adipose tissue was approved by the ethics committee of the University of Lübeck (EK Nr: 20-333), and the donor (male, 29 years) provided written consent.

## Supplementary

**Supplementary Figure 1:**
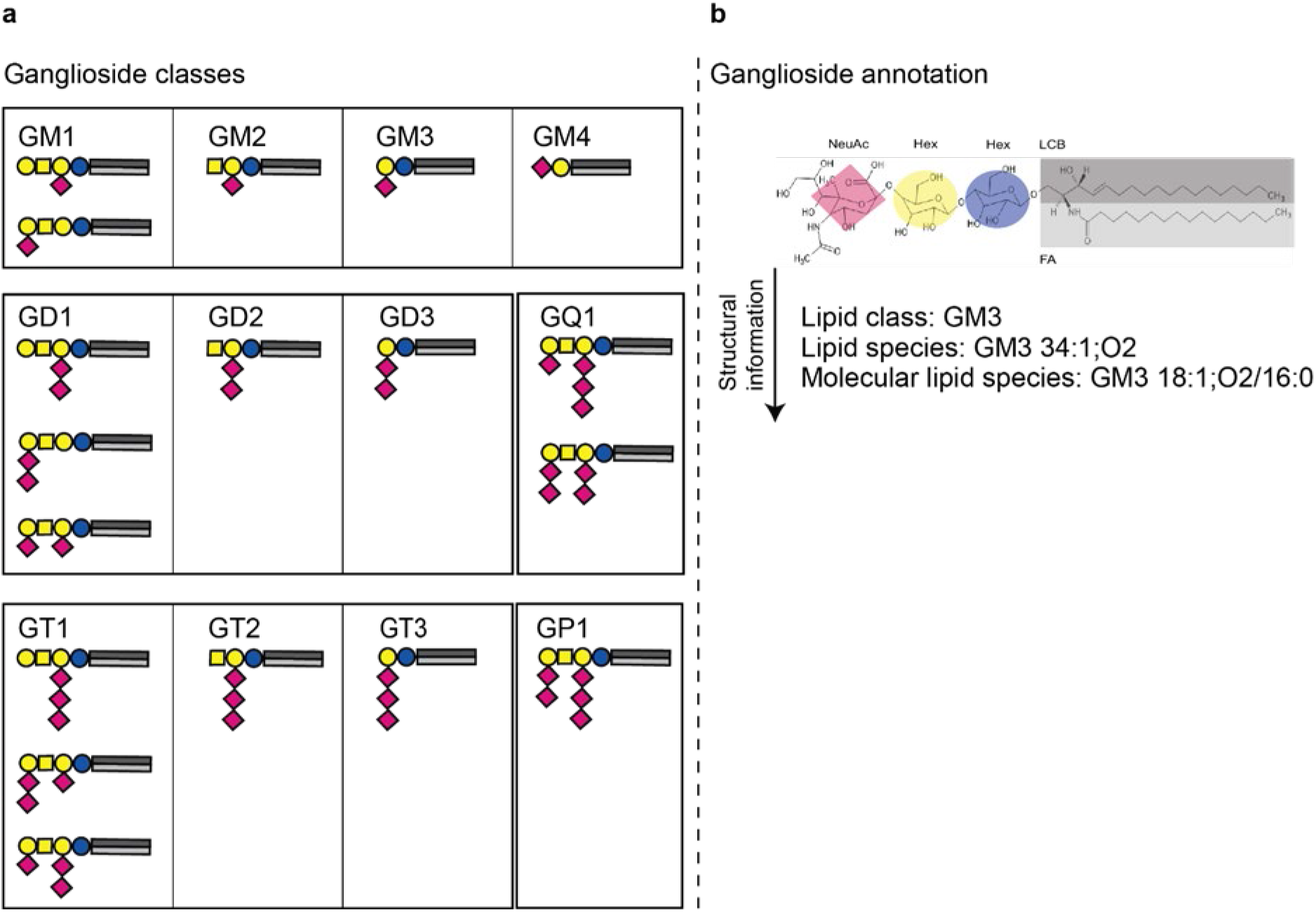
Ganglioside class and annotation overview. a Structures of the 12 different ganglioside classes containing one (GM1-4), two (GD1-3), three (GT1-3), four (PQ1) or five (GP1) sialic acid residues. b Ganglioside annotation based on the lipid class, species or molecular lipid species level.

**Supplementary Figure 2:**
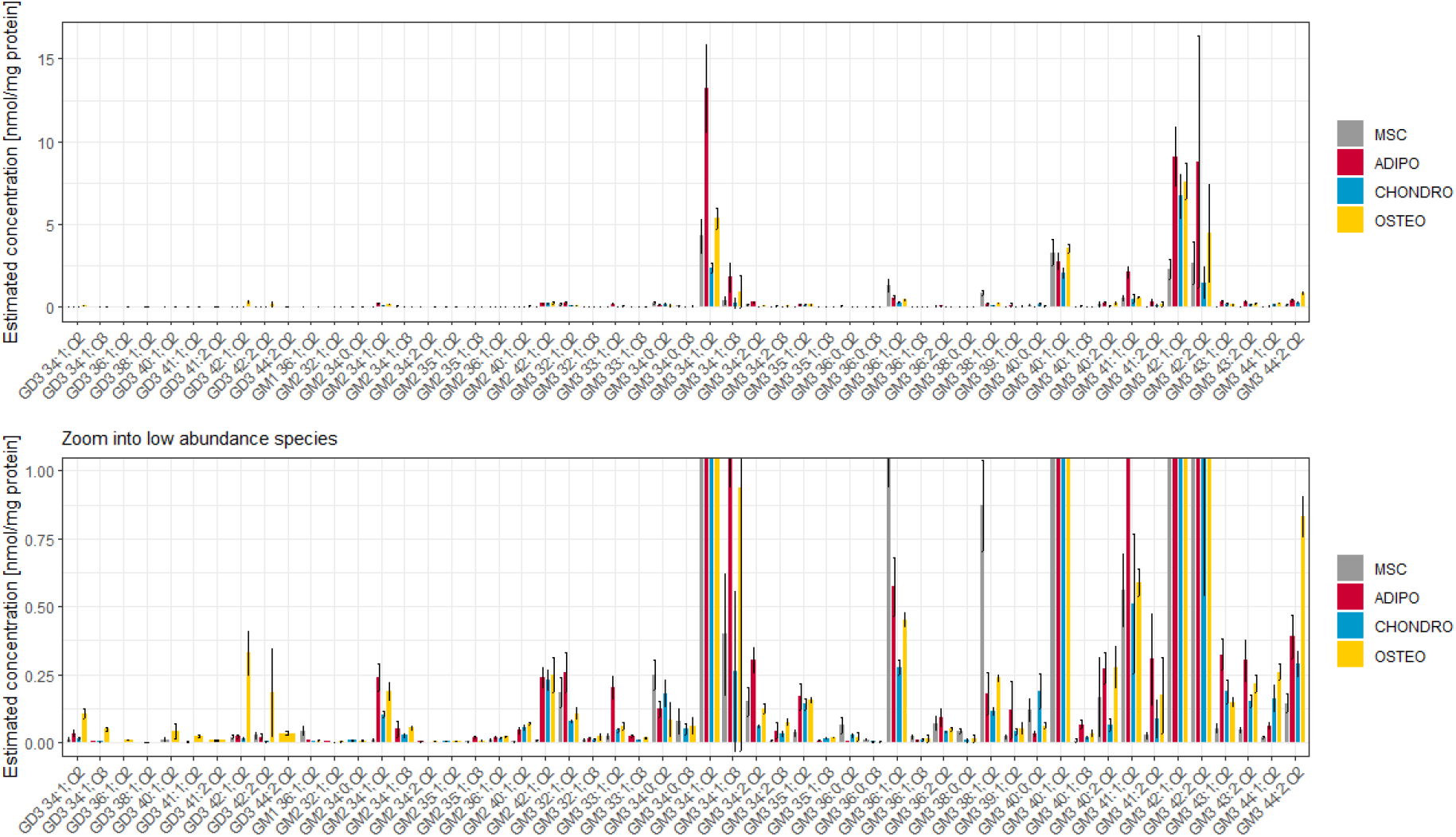
Overview on potential gangliosides markers in native and differentiated MSCs. Potential ganglioside marker on the lipid species level (adduct: [M-H]) including estimated concentration based on internal standardization with GM3 d5 18:1;O2/18:0 and normalization to the protein content.

**Supplementary Figure 3:**
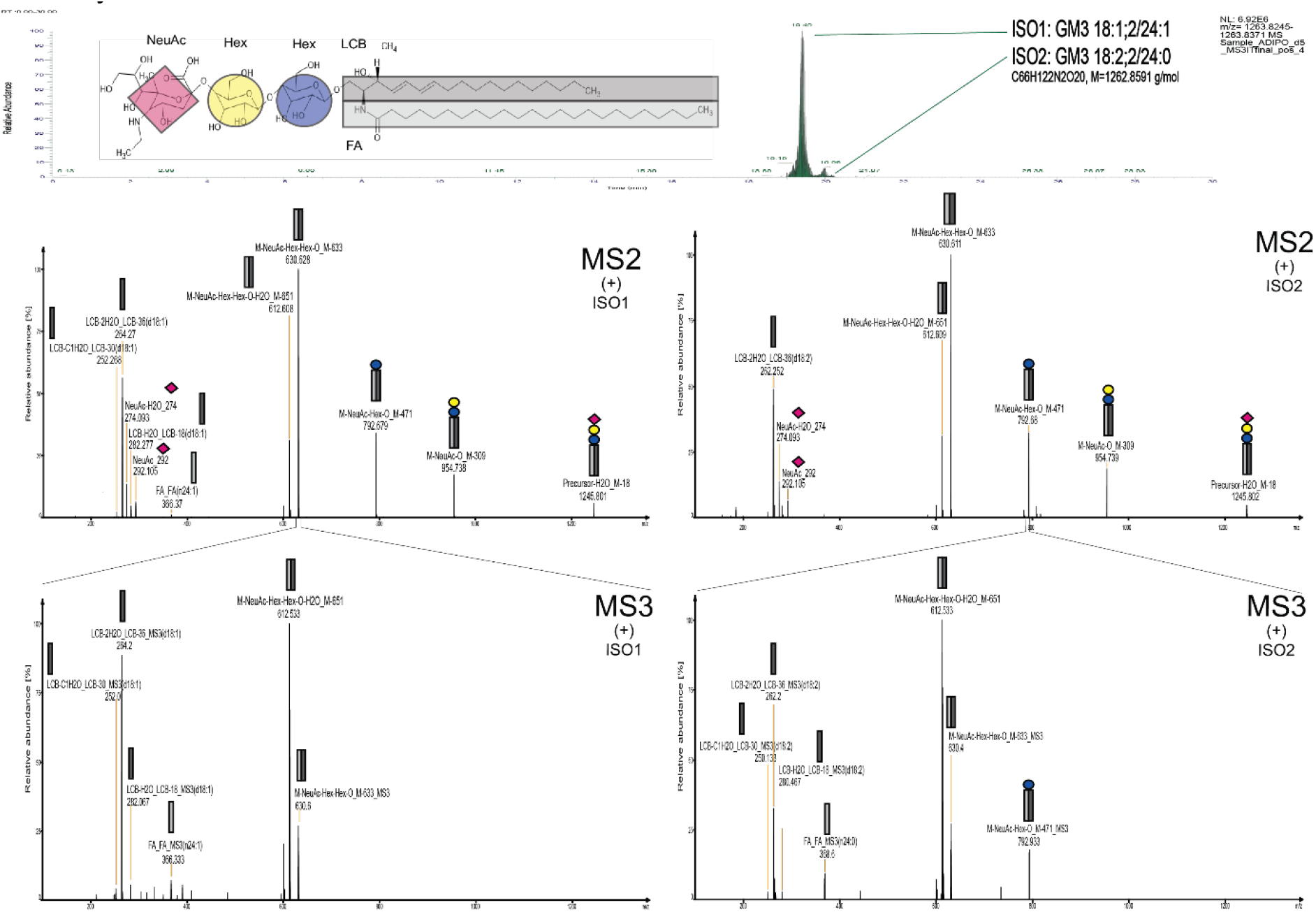
The benefit of combining reversed-phase chromatography and multistage fragmentation for ganglioside annotation. RP-HRMS^n^ enables isomeric separation of GM3 18:1;O2/24:1 at retention time 19.40 and GM3 18:2;O2/24:0 at retention time 19.96 followed by molecular species level assignment based on the fragmentation of the glycan and ceramide part in MS2 and MS3 (positive ion mode). GM3 18:2;O2/24:0 is an isomeric marker for adipocytes compared to GM3 18:2;O2/24:0 being upregulated in osteocytes.

**Supplementary Figure 4:**
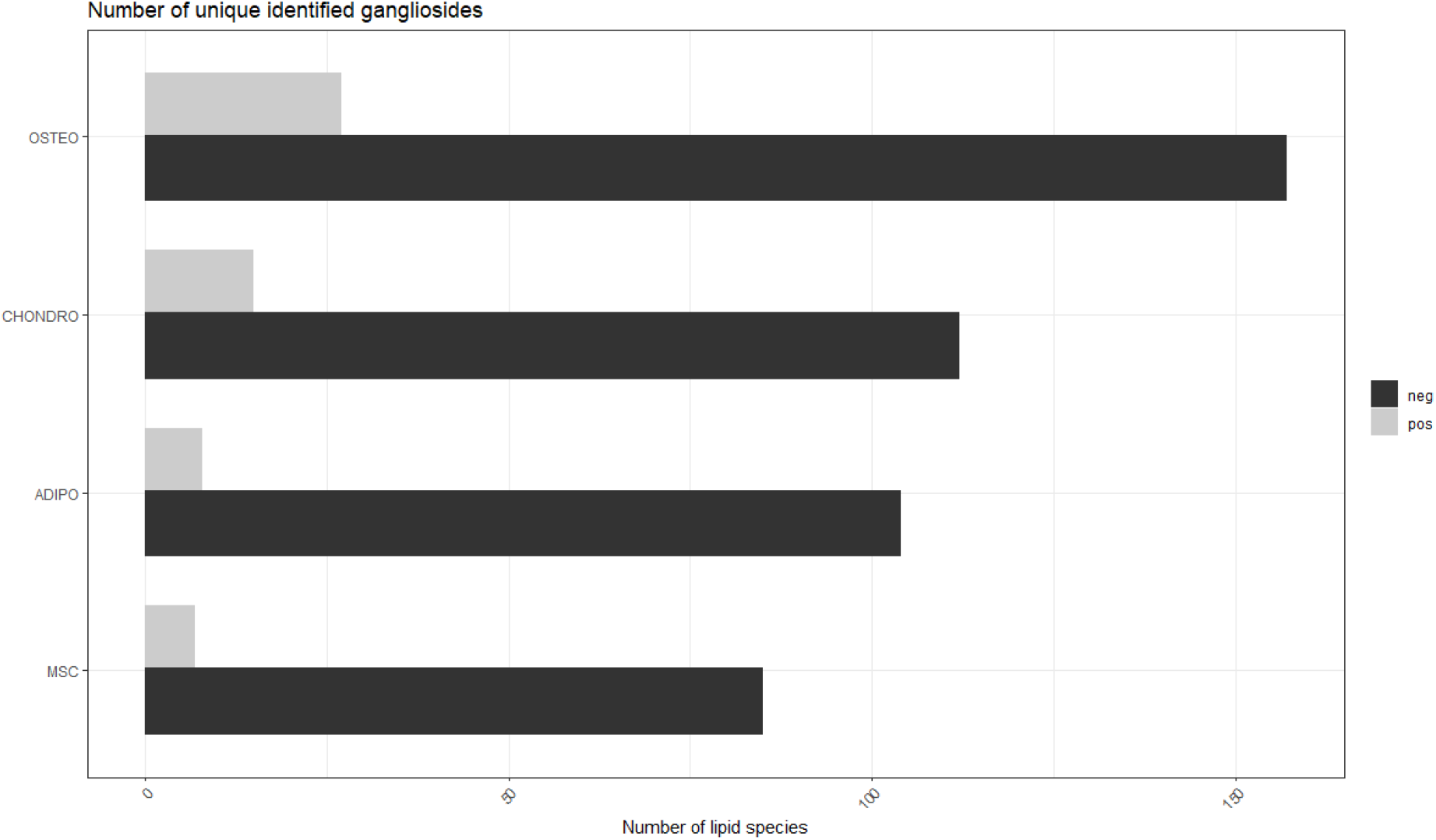
Gangliosides species numbers identified in the different cell states. An increasing number of ganglioside species were observed in all differentiated cells states (fat, bone, cartilage), with the highest number of gangliosides identified in osteocytes for both positive and negative ionization.

**Supplementary Figure 5:**
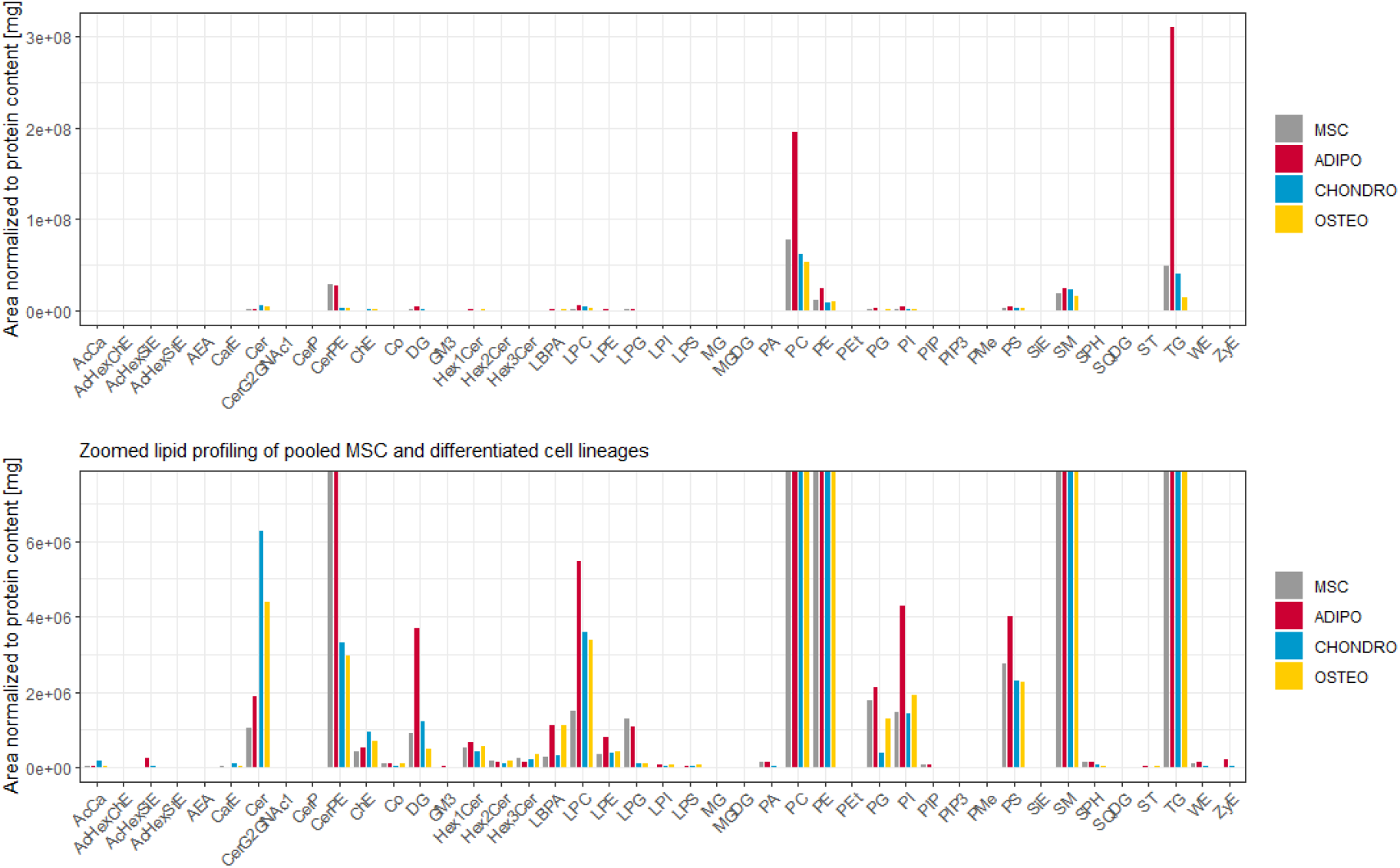
Lipid profiling of stem, fat, bone, and cartilage cell pools. For fat cells, TG, DG and PC are present in higher abundance. A potential cause may be the role of adipocytes acting as storage organelles for neutral lipids. Additionally, higher amounts of ceramides are detectable in cartilage (chondrocyte) cells.

**Supplementary Table 1: Overview of gangliosides marker**. List of annotated gangliosides (M-H adducts) in native and differentiated MSCs (n=5), including estimated concentrations based on normalization to GM3 and the protein content. The 57 identifications corresponding to the 16 previously unreported novel ganglioside molecular species are indicated by blue background. Species level annotation is provided in old (GM3 d36:1) and new (GM3 36:1;O2) nomenclature to avoid misinterpretation.

**Supplementary Table 2: Annotated gangliosides in native and differentiated MSCs**. List of annotated gangliosides (all adducts) in native and differentiated MSCs (n=5), including estimated concentrations based on normalization to GM3 and the protein content.

